# Egg cannibalism by passion vine specialist *Disonycha* Chevrolat beetles (Coleoptera: Chrysomelidae: Galerucinae: Alticini)

**DOI:** 10.1101/2020.04.15.005611

**Authors:** Colin R. Morrison, Wyatt Armstrong, Lawrence Gilbert

## Abstract

Cannibalistic behavior is now recognized to be an important component of nutritional ecology in both carnivorous and herbivorous species, including many beetle families (Englert and Thomas 1970; Beaver 1974; Dickinson 1992; Bartlett 1987; Alabi *et al.* 2008). This habit was historically viewed by an incidental outcome of unnaturally crowded laboratory situations with little ecological importance (Fox 1975), but it is increasingly acknowledged that cannibalism represents a potentially advantageous behavior (Richardson *et al.* 2010). Here we report on multiple cases of egg cannibalism, or conspecific oophagy, by adults of two species of passion vine (*Passiflora* Linnaeus: Passifloraceae) specialist flea beetles in the genus *Disonycha* Chevrolat (Coleoptera: Chrysomelidae: Galerucinae: Alticini). This is the first report of egg cannibalism from the Galerucinae, and to our knowledge, only the fourth report of egg cannibalism by adults in the Chrysomelidae; the other three reports are of adult Chrysomelinae species eating conspecific eggs (Dickinson 1992; McCauley 1992; Schrod *et al.* 1996). We conclude this note with several questions raised by our observations, followed by a discussion that may contribute to explanations of this behavior.

---

The genus *Disonycha* contains approximately 70 nominal species distributed widely from southern Canada to temperate South America and the Caribbean. Species richness is highest in northern temperate and subtropical latitudes, with the greatest number of species known from Mexico (Furth 2017). Adults are large for alticines (7 – 8 mm length), often with brightly colored and striped elytra. All are herbivorous as larvae and adults, with both life stages commonly inhabiting the same host plant. Three *Disonycha* species are known to specialize exclusively on *Passiflora*: *Disonycha discoida* Fabricius*, D. stenosticha* Schaeffer and *D. quinquelineata* Latreille. This note focuses on *D. stenosticha* and *D. quinquelineata* (Fig. 1). *Disonycha stenosticha* is distributed from the Rio Grande Valley of Southern Texas to Tamulipas and Veracruz states in south-central Mexico. Larvae and adults consume *P. suberosa* and *P. biflora* in the field, with unconfirmed reports of larvae consuming *P. fillipes* (Clark *et al.* 2004). *Disonycha quinquelineata* is found from the Mexican states of Guerrero, Veracruz and Yucatan to northwest Colombia. Larvae and adults eat *P. auriculata*, *P. biflora*, and newly flushed foliage of *P. pittieri*; adults will eat fresh *P. ambigua* leaves. Host records from the field have been confirmed by feeding trials in the laboratory, however they probably do not capture all host plants across these species’ ranges. Both species oviposit egg masses on the abaxial sides of host leaves, as well as leaves of non-host plants near larval foodplants (Fig. 1).

**Figure. 1.**
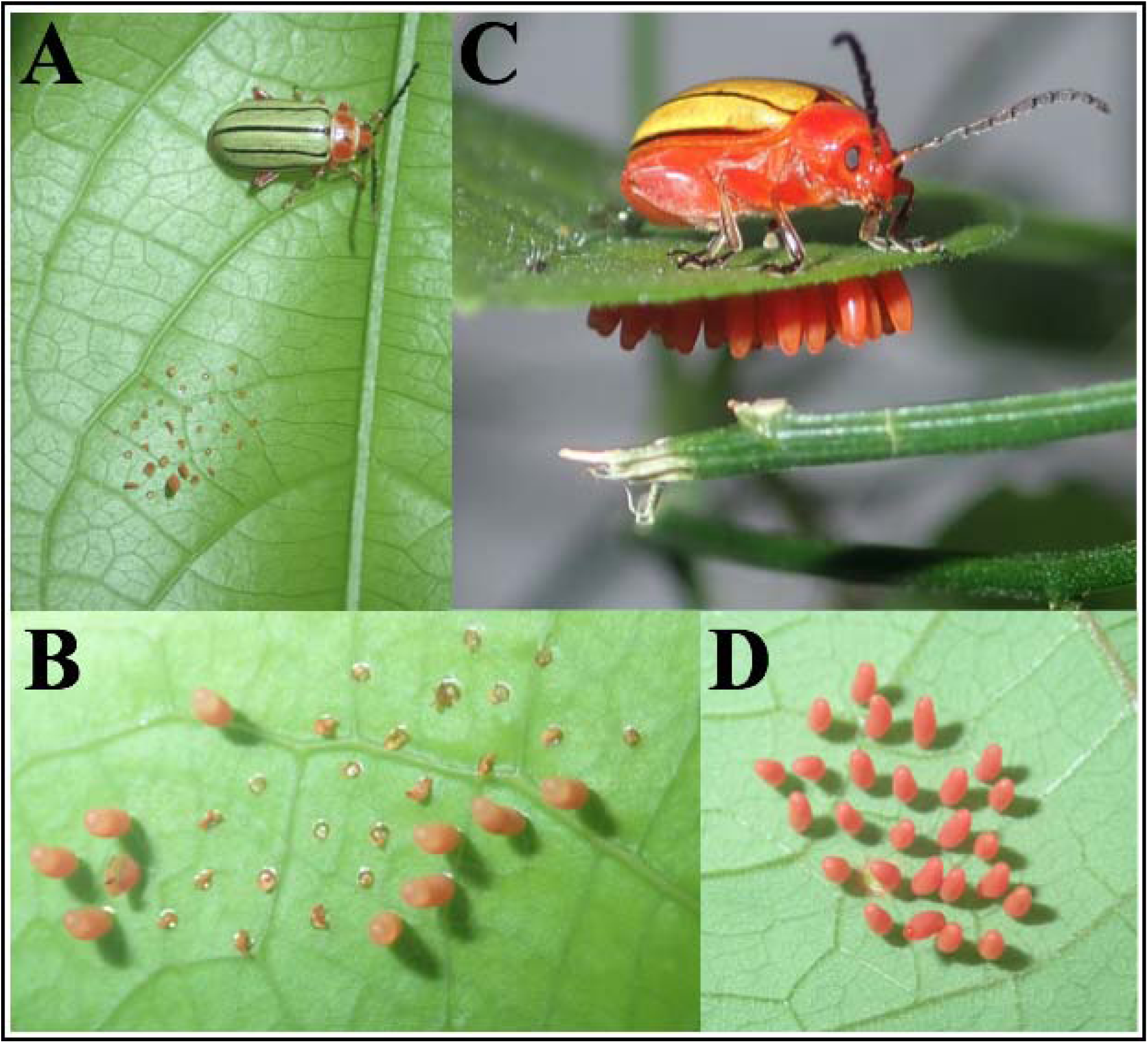
Passion vine specialist *Disonycha* egg cannibalism habits. A) Male *Disonycha quinquelineata* that just finished consuming all but one egg in a conspecific egg mass at La Selva Biological Station, Costa Rica; B) *D. quinquelineata* egg mass on *Passiflora ambigua* with 10/32 eggs consumed at La Selva; C) *D. stenosticha* female with newly oviposited egg mass on *P. suberosa* at National Butterfly Center, Mission, Texas; D) 3 d old *D. stenosticha* egg mass on *P. biflora* in research culture at UT Austin.

*Diconycha* egg cannibalism has been observed ten times, in both the field (N = 2) and the laboratory (N = 8). *Disonycha quinquelineata* was responsible for six of these events, and *D. stenosticha* for the other four. Egg masses have been consumed by both males (*D. stenosticha* N = 2 individuals, *D. quinquelineata* N = 2) and females (*D. stenosticha* N = 2 individuals, *D. quinquelineata* N = 4), including two instances where an individual *D. quinquelineata* female consumed her own eggs within one day of oviposition. All adult egg cannibalism incidents occurred within 36 h of oviposition. This is about how much time it takes for the chorion to harden to the point where it is stiff to the touch. Prior to this, the eggs are soft and palatable. In three cases the adult did not eat all the eggs in each mass; 65 – 70 % of the eggs were consumed in each of these instances (Fig. 1). *Disonycha quinquelineata* and *D. stenosticha* neonate larvae will also cannibalize unhatched eggs from the same clutch shortly after eclosing. It is not known if all eggs that are consumed by neonates are viable, or unfertilized trophic eggs oviposited to provision the first larvae to eclose (*e.g.* Perry and Roitberg 2006). It is not within the scope of this contribution to report on *Disonycha* larval cannibalism, however it is important to the context of this observation to note that this also occurs.

Cannibalistic juvenile behavior is relatively common among generally non-carnivorous insect orders (78% percent of 178 studies reviewed by Richardson *et al.* 2010). However, adult consumption of conspecific eggs is less common, less frequently observed, or underreported. Adult egg cannibalism has been noted in the Blattodea, Hemiptera, Hymenoptera, Orthoptera, and Coleoptera. Beetle families where adult consumption of conspecific eggs has been observed include: Zopheridae (Kubik *unpublished data*), Tenebrionidae, Coccinellidae, Cucujidae, Dermestidae, Scolytidae, Silphidae, and Chrysomelidae (*see references within* Richardson *et al.* 2010). Two subspecies of adult milkweed leaf beetles, *Labidomera c. clivicollis* Kirby and *L. c. rogersii* LeConte, will consume conspecific egg masses (Dickinson 1992). Conspecific egg cannibalism has also been observed in the Colorado potato beetle, *Leptinotarsa decemlineata* Say (Schrod *et al.* 1996), and the false Colorado potato beetle, *L. juncta* Germar (McCauley 1992). Cases of egg cannibalism by larvae are more commonly reported for chrysomelid beetles (*e.g.* Eberhard 1981; Windsor *et al.* 2013). Chrysomelid larvae are also known to eat unfertilized trophic eggs. However, this is not egg cannibalism *sensu stricto*, in that these unviable eggs are intentionally laid for purposes other than producing offspring directly. For example, gravid *L. decemlineata* mothers experiencing predation risk from stinkbugs lay unfertilized trophic eggs to supplement the nutrition of their offspring (Tigreros *et al.* 2017). Upon hatching, neonate larvae consume the trophic eggs, giving them extra nutrition to grow quickly while avoiding exposure to stink bugs, and other natural enemies, while foraging on leaves. Egg viability was not assessed in all previous reports of chrysomelid larvae consuming conspecific eggs. Thus, prevalence of true egg cannibalism by larvae versus offspring resource provisioning with trophic eggs remains unclear.

It is important to disentangle the social and ecological contexts under which beetle cannibalism occurs (Trumbo and Valletta 2007; Wood *et al.* 2014). Egg cannibalism is generally attributed to density-dependent factors, such as crowding situations in which high larval abundances lead to rapid consumption of food resources that result in starvation of the whole group. In a review of insect cannibalism, Richardson and colleagues (2010) attributed consumption of beetle conspecifics to density-dependent factors in 70% of studies reporting cannibalism. However, density-dependent resource scarcity alone does not account for the frequency of egg cannibalism by *Disonycha* adults. Cannibalized egg masses were consumed in locations with abundant host plant material in both the laboratory and the field, in all cases. Two instances of a *D. quinquelineata* adult consuming eggs in the field occurred at the La Selva Biological Reserve in Costa Rica (10°25’53” N / 84°00’17” W) in a large *Passiflora* garden (~ 15 x 15 m) with copious foodplant for adults to eat. In the laboratory at The University of Texas at Austin, egg masses were consumed in large (0.53 m^3^), medium (0.07 m^3^), and small (0.03 m^3^) mesh pop-up cages replete with fresh *Passiflora* shoots or potted host plants with abundant new growth. Cannibalism occurred in cages where egg masses were present on nearby foodplants (N = 4), as well as in cages without eggs (N = 4). Egg consumption occurred in cages with different densities and adult sex ratios. Specifically, cannibalism occurred in cages housing multiple adults of both sexes (N = 4), cages with a single mated pair (N = 2), and cages with only the female that oviposited the eggs (N = 2). This is evidence for the hypothesis that relative lifestage abundance is not linked to egg cannibalism. The influence of larval presence on adult egg cannibalism cannot be inferred because eggs, adults, and larvae were never housed together in the same cage.

Density-independent factors such as poor host plant nutritional quality (O’Rourke and Hutchinson 2004), and developmental asynchrony within populations (Nakamura and Ohgushi 1981) can also influence herbivorous insect cannibalistic behavior. Investigation of the relative influence of environmental and demographic variables on frequency of egg cannibalism by *Passiflora* specialist *Disonycha* flea beetles is open for study.

Newly oviposited eggs may represent a concentrated nutritional supplement to beetles that may not be receiving sufficient protein from host plant material alone. *Disonycha stenosticha* and *D. quinquelineata* eggs have higher protein concentration per unit mass than new leaves of the principal host plants that they feed on in the field (Fig. 2). This stoichiometric discrepancy is true of individual eggs, and the entire egg mass relative to foodplant material. Protein concentration comparisons were made between *D. quinquelineata* eggs and *P. biflora* (t-test; N = 8; *P* < 0.005), *D. stenosticha* eggs and *P. suberosa* (t-test; N = 5; *P* < 0.001). These data support the hypothesis that consumption of one’s own eggs may arise from a need to re-assimilate protein stores necessary for maximizing performance, and by extension their fitness. One possibility is that consuming egg masses with a disproportionate protein concentration relative to host leaves would improve a *Disonycha* beetle’s current chances of surviving so that reproductive effort can be maximized in the future. Such a situation has been described with cannibalistic *Coleomegilla maculata lengi* De Geer larvae (Coleoptera: Coccinellidae) that displayed strong preference for eating conspecific eggs over aphid prey (Gagné *et al.* 2002); preference for egg consumption correlated with superior performance leading the authors to infer that *C. m. lengi* eggs had higher nutritional quality than equivalent quantities of aphids.

**Figure. 2.**
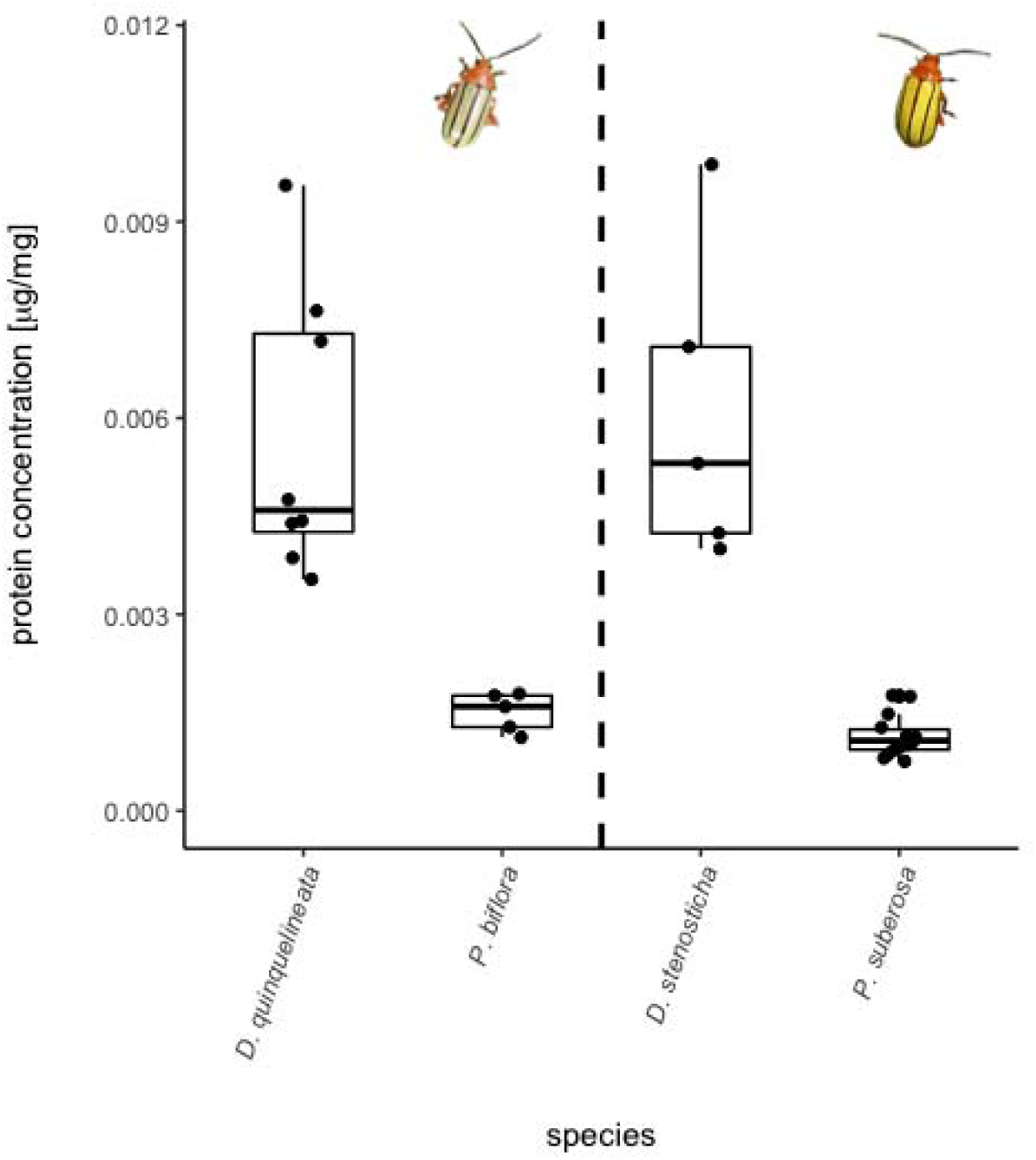
Comparison of *Disonycha quinquelineata* and *D. stenosticha* egg mass and host plant leaf protein concentration. Beetle species are paired with their preferred host plants on each side of the vertical dashed line. These data were analyzed with Welch’s t-tests.

*Disonycha stenosticha* and *D. quinquelineata* adults frequently live for six to nine months in the laboratory (we assume at least half this long in the field), and females can oviposit multiple egg masses over this period. These long-lived adults have multiple opportunities to pass on their genes. However, different selection pressures create costs on gamete production and resource investment that differ for males and females (Blum and Blum 2012). Males are expected to mate with as many females as possible. If a male has mated with multiple females, cannibalism of a given egg mass represents a low risk of consuming his own offspring. Cannibalism of egg masses oviposited by other females would be advantageous to the relative fitness of a female that is still sourcing her offspring into the population. Consuming her own eggs does not appear beneficial because she would be decreasing her relative fitness in that population. However, there exist external constraints on female reproductive investment and survival that were not accounted for here (*e.g.* living to produce future clutches of healthy offspring) that prevent us from making strong inferences about what would be selectively advantageous to a beetle that eats her own eggs.

Egg cannibalism by conspecifics may contribute to the difficulty finding high density populations of *Disonycha* in the field. With the exception of semi-artificial habitats such as native plant nurseries in South Texas and passion vine gardens in Costa Rica, we have not observed *Disonycha* beetles aggregating at host plant patches in high numbers. Larval *Disonycha* consume a greater amount of plant material per unit time than adults and the *Passiflora* that host these species are usually not abundant in the habitats where they occur. *Passiflora* primarily occur in spatially and temporally ephemeral gaps, and edge habitats where competition for space and light is intense. This creates a situation where populations that are not self-regulated and/or diminished in size by interspecific competition and mortality from natural enemies could easily defoliate host plant patches resulting in resource scarcity, potentially resulting in local starvation. Egg cannibalism may serve as a population level mechanism to dampen the number of individuals competing for these scarce resources.

It is possible that *Disonycha* adults are only consuming unfertilized eggs. We considered this caveat for three reasons: 1) adults did not always consume every egg in a mass; 2) *D. quinquelineata* egg mass dissections (N = 10) showed a range of egg viability ranging from 0 to 95%; and 3) no *D. stenosticha* or *D. quinquelineata* egg masses in which any larvae eclosed have had 100% hatching success. Despite this caution, we reason that specific consumption of unfertilized eggs is not likely to be the case. Fertilized and unfertilized eggs appeared to be randomly distributed in a given egg mass, and adults proceeded to eat eggs indiscriminately from the periphery of the egg mass to a specific point where consumption ceased during events in which less than 100% of eggs were consumed. Thus, egg cannibalism is occurring unless *Diconycha* beetles have the capacity to discriminate between viable and unviable eggs (*e.g.* homogenous egg dispersal in bruchid beetles; Messina *et al.* 1987), or they are only consuming masses that contain unviable eggs.

The observations reported here suggest that egg cannibalism by *D. stenosticha* and *D. quinquelineata* adults is meaningful for the fitness of these species. These are the main questions that these observations compelled us to ask: 1.) Does genetic relatedness between eggs and adults affect an individual’s propensity to cannibalize eggs? 2.) Is egg cannibalism density-dependent? 3.) Are trade-offs in propensity to eat eggs different over the course of an adult’s life? 4.) Are adults only consuming unfertilized eggs? 5.) Is egg cannibalism an important determinant of *Disonycha* population structure within the greater *Passiflora* specialized herbivorous insect community? The observations presented here provide a foundation for turning these questions into testable hypotheses. Specifically, we hypothesize that genetic relatedness does not predict propensity to cannibalize eggs as we have observed females eating their own egg masses as well as those of other individuals. Egg cannibalism occurs regardless of egg mass density in a given area but occurs more frequently at higher egg mass densities. Eggs provide adults with more protein than equivalent amounts of host leaves and this disparity drives age-specific variation in cannibalism rates. These beetles are long-lived, and with such longevity comes the eventuality of protein limitation for an individual to generate new gametes and maintain cell division and repair processes (*e.g.* Dunlap-Pianka et al. 1977). Thus, we predict older adults that have used up protein stores from the immature stage will cannibalize eggs more often than younger adults. Adults eat eggs regardless of whether they are viable. Discrimination between viable and unviable eggs is unlikely given that viable eggs are oviposited among unfertilized eggs in a given mass. However, the ability to differentiate between viable and unfertilized eggs via chemosensory pathways could provide for such discriminatory ability. Such pathways have been reported on in the Chrysomelidae (Mitchel 1994; van Loon 1996; Fernandez and Hilker 2007). Many proximate factors contribute to variation in population density and host usage by *Disonycha* and other members of the *Passiflora* specialist insect community (Smiley 1982; Gilbert 1991; Engler and Gilbert 2007). Egg cannibalism may contribute to this heterogeneity by maintaining the relatively uniform spatial distribution of *D. stenosticha* and *D. quinquelineata* individuals that we have observed in nature.

Cannibalism can be a significant driver of an organism’s life history, ecology, and fitness. Unfortunately, this behavior is underreported, and studied empirically even less frequently. There is reason to think that adult insects functionally characterized as herbivores cannibalize conspecific eggs more often than what the current literature currently describes. This is a rich area for study, and we encourage researchers with knowledge of their study organisms to report on this behavior when the opportunity is presented.

## Acknowledgments

Many thanks to John Smiley for opening the door to the fascinating world of *Passiflora* specialized flea beetles and supporting us to investigate diverse aspects of the system. Thoughtful critique of earlier versions of the manuscript were provided by T. Kubik. Funding to support this research was provided by the Texas Ecological Laboratories Program and the Organization for Tropical Studies, Costa Rica.

## Notes

### Competing Interest Statement

The authors have declared no competing interest.

